# Enhanced translational activity is linked to lymphatic endothelial cell activation in cutaneous leishmaniasis

**DOI:** 10.1101/2024.08.07.605632

**Authors:** Lucy Fry, Hayden Roys, Anne Bowlin, Gopinath Venugopal, Jordan T. Bird, Alexx Weaver, Stephanie D. Byrum, Tiffany Weinkopff

## Abstract

Cutaneous leishmaniasis (CL) is a significant public health problem leading to permanently disfiguring skin lesions caused by *Leishmania* parasites. Lesion severity stems from an excessive host inflammatory response that prevents healing. Here, we characterized the transcriptional and translational responses of lymphatic endothelial cells (LECs) during murine CL using historical single-cell RNA sequencing data combined with flow cytometry and in vivo puromycin incorporation to assess translational activity. We identified upregulation of antigen presentation pathways including MHC-I, MHC-II, and immunoproteasome transcripts in dermal LECs from *Leishmania major*-infected mice compared to naive controls. LECs also exhibited increased expression of guanylate binding proteins and interferon-inducible genes, indicative of immune activation. Moreover, our findings demonstrate that LECs in leishmanial lesions displayed heightened translational activity relative to LECs from uninflamed ears, and LEC translational activity was highest in activated LECs. Furthermore, LEC translational activity exceeded that of other cell types within the lesion microenvironment. Validating the transcriptomic data, LECs in lesions expressed elevated MHC-II and programmed death-ligand 1 (PDL-1), supporting their potential role in antigen presentation. Functional assays using DQ-OVA confirmed that LECs from leishmanial lesions efficiently uptake and process antigens, highlighting their capability as antigen presenting cells in the inflamed dermal microenvironment. Overall, our study reveals the activation status of LECs in leishmanial lesions, shedding light on their potential role in shaping local immunity and inflammation in a variety of skin diseases.

## INTRODUCTION

Cutaneous leishmaniasis (CL) is a significant public health problem caused by *Leishmania* parasites. Characterized by permanently disfiguring, ulcerative skin lesions, CL is the most common clinical form of leishmaniasis, with ∼ 1.5 million new cases annually and 12 million people infected worldwide (1). No CL vaccine exists, and available therapies are often ineffective without multiple rounds of treatment (2). Lesion severity is driven by an excessive inflammatory response that prevents healing. Large numbers of circulating immune cells, such as inflammatory monocytes and neutrophils, are recruited to the infection site, promoting host immunopathology and exaggerating inflammation (3–6). Parasites are skin tropic and largely restricted to the lesion site without extensive dissemination. Specifically, dermal macrophages harbor the parasites. Parasites are controlled by CD4^+^ T cell-produced IFN-γ. Notably, even when parasite burdens are controlled, cutaneous lesions often persist, suggesting other factors—such as the host inflammatory response—contribute to disease pathogenesis (7–12). Due to persistent CL, there is a critical need to develop new host-directed therapeutic strategies that reduce inflammation and promote healing.

Lymphatic remodeling (changes in lymphatic endothelial cells (LECs) or lymphatic vessels (LVs)), including lymphangiogenesis (formation of new lymphatic vessels [LVs]) and lymphangiectasia (enhanced LV dilation), occurs during inflammation and dictates lymphatic function. We found that lymphatic remodeling takes place at the dermal lesion after inoculation with *Leishmania major* (13–15). The frequency of LEC proliferation is higher in leishmanial lesions than uninfected skin, suggesting enhanced lymphangiogenesis (15). Mimicking human leishmanial lesions, the transcription factor HIF-1α (hypoxia-inducible factor-1α), its target VEGF-A (vascular endothelial growth factor-A), and the VEGF-A receptor VEGFR-2, are elevated in lesions during murine leishmaniasis (14–18). Furthermore, blocking VEGFR-2 signaling impairs LEC proliferation and increases immunopathology with no effect on parasite load (15). We identified macrophages as the major cellular source of VEGF-A in lesions and showed that VEGF-A expression requires myeloid aryl hydrocarbon receptor nuclear translocator (ARNT)/HIF-α signaling (14). Importantly, myeloid ARNT/HIF-α signaling promotes lymphangiogenesis to limit immunopathology, suggesting lymphangiogenesis is required for healing (13, 19). Therefore, these data suggest that dermal lymphangiogenesis is required to restrict immunopathology and resolve dermal lesions.

Because lymphangiogenesis attenuates CL severity, we wanted to further characterize LECs in leishmanial lesions. In a previous studying subjecting murine leishmanial lesions to scRNASeq, we found LECs from infected ears expressed increased guanylate binding proteins (*Gbp2, Gbp4,* and *Gbp7*) and a higher abundance of both MHCI and MHCII transcripts (*H2-K1, H2-D1, H2-Q7, H2-Aa, H2-Ab1*) (20).

*Cxcl9* was significantly elevated in LECs following *L. major* infection. Following Ingenuity Pathway Analysis (IPA) comparing LECs from infected skin and uninfected control skin, we identified predicted upstream regulators of LECs including *IFNγ* inducing activation, while *IL-10* downregulates LEC activation upon infection (20).

Predicted transcriptional regulators activating LECs include *IRF3, STAT1,* and *IRF7*, and predicted transcriptional regulators inhibiting LECs include *MLXIPL, MYC, SIRT1, TRIM24*, and MYCL (20). Based on elevated transcripts *B2m* and *H2-*K1, IPA analysis also revealed the antigen presentation and PD1/PD-L1 pathways are increased with infection in LECs (20). In contrast, IPA analyses showed LECs from infected mice exhibit decreased EIF2, mTOR, and eIF4/p70S6K signalling, hinting at an important role for oxidative stress at the site of infection (20). These transcriptomic findings indicate that in addition to their function in draining fluid and cells away from the site of infection, dermal LECs participate in the host immune response during CL.

Here, we validate our scRNASeq findings and show that LECs are activated exhibiting a mixed phenotype, characterized by pro-inflammatory molecules such as GBPs and anti-inflammatory co-stimulatory molecules like PDL-1, during *L. major* infection in vivo. Our findings contrast with the widely accepted current view that LECs mainly exhibit an immunosuppressive phenotype (21–30). We also find LECs exhibit enhanced translational activity that is linked to LEC activation. These results suggest that increases in LEC numbers due to proliferation not only help resolve lesions via their vessel drainage functions, but that LECs could participate in anti-parasitic immunity or restricting parasite dissemination. These findings demonstrate for the first time that LEC activation is a feature of the pathogenesis of CL.

## MATERIALS AND METHODS

### Mice

Female C57BL/6NCr mice were purchased from the National Cancer Institute. Mice were housed in the Division of Laboratory Animal Medicine at University of Arkansas for Medical Sciences (UAMS) under pathogen-free conditions. Mice between 6 and 8 weeks of age were used for infections. All procedures were performed in accordance with the guidelines of the UAMS Institutional Animal Care and Use Committee (IACUC).

### Parasites and infections

*Leishmania major* (WHO/MHOM/IL/80/Friedlin) parasites were maintained in vitro in Schneider’s Drosophila medium (Gibco) supplemented with 20% heat-inactivated FBS (Invitrogen), 2 mM L-glutamine (Sigma), 100 U/mL penicillin, and 100 mg/mL streptomycin (Sigma). Ficoll density gradient separation was used to isolate metacyclic stationary phase promastigotes from 4–5 day cultures by (Sigma) (31). For dermal ear infections in mice, 2×10^6^ promastigote parasites in 10 µL PBS (Gibco) were injected intradermally into the ear. For analyses, ears were excised, dorsal and ventral sheet were separated. Ear sheets were enzymatically digested for 90 min at 37°C using 0.25 mg/mL Liberase (Roche) and 10 mg/mL DNase I (Sigma) in incomplete RPMI 1640 (Gibco). After digestion, ears were smashed through a filter to obtain a single-cell suspension (20).

### Flow cytometry

To assess cell viability, cells from both infected and contralateral ears were incubated with fixable Aqua dye (Invitrogen) or Zombie (Biolegend) for 10 min at room temperature. Cells were treated with FcγR blocking reagent (Bio X Cell) and 0.2% rat IgG for 10 min at 4° before staining for the following markers: anti-CD4-PE (GK1.5) and anti-CD11c-PE (N418) were purchased from eBioscience; anti-Ly6C-PerCP-Cy5.5 (clone HK1.4), anti-Ly6G-eFlour 450 (clone 1A8), anti-CD8β-eF780 (H35-17.2), anti-CD31-PECy7 (390), and anti-CD45-AF700 (clone 30-F11), were purchased from Invitrogen; anti-CD64-BV711 (clone X54-5/7.1), anti-CD11b-BV605 (clone M1/70), anti-CD31-AF488 (390), anti-MHCII-APCeF80 (M5/114.15.2), anti-PDL1-BV650 (10F.9G2) and anti-podoplanin-PE/Dazzle 594 (clone 8.1.1) were purchased from BioLegend; anti-CD24-BV560 (M1/69) was purchased from BD Biosciences. Surface staining was performed in Brilliant Violet Buffer (BD Biosciences) or Super Bright staining buffer (eBiosciences). Intracellular cytokine staining was performed using the Foxp3 intracellular staining kit (Life Technologies) combined with α-CD3 APCeF780 (Invitrogen). Cells were acquired using an LSRII Fortessa (BD Biosciences) or Northern Lights (Cytek) flow cytometers and analyzed using FlowJo software version 10.2 (Tree Star).

### In vivo translation

To determine the translational activity of LECs in vivo, C57BL/6 mice were infected, and translation was measured by puromycin incorporation using flow cytometry as previously described (32, 33). In brief, mice are injected with 21.8 μg/g of puromycin (Sigma P7255) diluted in PBS at 1 hour prior to euthanization. Puromycin was detected by flow cytometry using an anti-puromycin antibody conjugated to A647 (Sigma MABE343-AF647) after intracellular staining with the Foxp3 kit (Life Technologies).

### DQ-OVA antigen uptake and presentation

DQ ovalbumin (DQ-OVA) (Invitrogen D-12053) is labeled with a pH-insensitive, green fluorescent Bodipy FL dye. For ex vivo assays, ears were digested, processed, and counted. A single cell suspension of ear cells in 500 μL of complete RPMI were cultured in 5 mL Falcon polypropylene tubes at 37°. For antigen uptake assays, cells were cultured with DQ-OVA (10 μg/mL) for 10 min. For antigen processing assays, cells were cultured with DQ-OVA (10 μg/mL) for 2 hours. DQ-OVA uptake and processing was analyzed by flow cytometry in the Af488/FITC channel. Cells cultured without DQ-OVA for 2 hours provided FMO controls. For antigen presentation assays in vivo, DQ-OVA (20 μg/lesion) was intradermally injected into leishmanial lesions in 20 μL of PBS. After 24 hours, ears were harvested, digested, processed, and stained for flow cytometry to examine DQ-OVA MFI in individual cell populations.

### scRNASeq samples preparation

The scRNASeq samples were prepared and used as a part of a previous study (20). In brief, the Arkansas Children’s Research Institute (ACRI) Genomics and Bioinformatics Core prepared NGS libraries from fresh single-cell suspensions using the 10X Genomics NextGEM 3’ assay for sequencing on the NextSeq 500 platform using Illumina SBS reagents. Trypan Blue exclusion under 10X magnification determined cell quantity and viability. Library quality was evaluated with the Advanced Analytical Fragment Analyzer (Agilent) and Qubit (Life Technologies) instruments.

### scRNASeq data analysis

Data analysis was carried out as a part of a previous study (20). Briefly, the UAMS Genomics Core generated Demultiplexed fastq files which were analyzed using 10X Genomics Cell Ranger alignment and gene counting software, a self-contained scRNASeq pipeline developed by 10X Genomics. The reads were aligned to the mm10 reference transcriptomes using STAR and transcript counts were generated (34, 35).

The *Seurat* R package was used to process the raw counts generated by *cellranger count* (36, 37). Potential doublets, low quality cells, and cells with a high percentage of mitochondrial genes were filtered out. Cells that have unique feature counts > 75^th^ percentile plus 1.5 times the interquartile range (IQR) or < 25^th^ percentile minus 1.5 time the IQR were filtered. Similarly, cells with mitochondrial counts falling outside the same range for mitochondrial gene percentage were filtered. After filtering, all 8 sequencing runs were merged. The counts were normalized using the LogNormalize method which log-transforms the results (20). Subsequently, the 2000 highest variable features were selected. The data was scaled, and Principal component analysis (PCA) was performed. A JackStraw procedure was implemented to define the significant PCA components that have a strong enrichment of low p-value features.

A graph-based clustering strategy was used to embed cells in graph structure (38). Seurat was used to visualize the results in t-distributed stochastic neighbor embedding (tSNE) and Uniform Manifold Approximation and Projection (UMAP) plots (39). Seurat *FindNeighbors* and *FindClusters* functions were optimized to label clusters.

Seurat *FindAllMarkers* function finds markers that identify clusters by differential expression, defining positive markers of a single cluster compared to all other cells and comparing those to known markers of expected cell types from previous single-cell transcriptome studies. Cell type determinations were decided by manually reviewing these results, and some clusters were combined if their expression was found to be similar. We specifically examined the Prox-1^+^ LEC cluster. For the heat maps showing individual differentially expressed genes (DEGs) within a specific pathway in the LEC cluster, the expression log ratio is reported.

### Statistics

All data were analyzed using GraphPad Prism 9. A Grubbs’ test was used to identify and mathematically remove outlier data points. Statistical significance was calculated using an unpaired 2-tailed Student’s t-test for a single comparison between groups. Statistical significance was also calculated using two-way ANOVA with a Tukey’s multiple corrections for multiple comparisons between groups and cell types. The test used is listed in the figure legend. For either test, *p* ≤ 0.05 was considered statistically significant.

### Artificial Intelligence or Large Language Models Statement

During the preparation of the text for this work throughout July 2024, the authors used ChatGPT 40 mini to improve readability and language. After using this tool, the authors confirm that they have reviewed and edited the content as needed, and they take full responsibility for the content of the publication.

## RESULTS

To define the transcriptional signature of LECs during cutaneous leishmaniasis, we infected C57BL/6 mice in the ear dermis with *Leishmania major* parasites and performed scRNASeq on leishmanial lesions as part of a previous study (20).

Specifically, we analyzed 13,034 total cells from naive ears and 13,524 cells from infected ears (20). Among the total cells, here we focus on the LEC cluster. IPA analysis revealed the ‘antigen presentation’ and ‘PD1/PD-L1’ pathways are increased with infection in LECs compared to LECs from naïve uninfected, uninflamed skin (20). Upon deeper analysis of the LEC transcriptome after dermal *L. major* infection, we find a higher abundance of beta-2 microglobulin and both MHC-I and MHC-II transcripts (*H2-K1, H2-D1, H2-Q7, H2-Aa, H2-Ab1, H2-Eb1, H2-Q4, H2-Q6, H2-T2*) in LECs with infection compared to naïve controls (Fig. 1A). Similarly, we also detect a higher abundance of the invariant chain (*Cd74*) and immunoproteasome transcripts (*Psmb8*, *Psmb9,* and *Psme2*) in LECs with infection compared to controls (Fig. 1A). We discovered LECs from infected ears expressed increased guanylate binding proteins (*Gbp2, Gbp3, Gbp4, Gbp5, Gbp6, Gbp7, Gbp8,* and *Gbp9*) and transcripts of interferon-inducible genes (*Cxcl9*, *Ifi44*, *Igtp*, *Iigp1, Irgm1*, and *Irgm2*) (Fig. 1B) (20). Based on elevated *Cd274* transcript (Fig. 1B), IPA analysis also revealed the ‘PD1/PD-L1’ pathway is increased with infection in LECs (20). This deeper investigation of our transcriptomic findings suggest that while parasites do not live inside LECs (20), LECs are becoming activated by responding to the inflamed tissue microenvironment at the site of infection by increasing their abundance of pro-inflammatory transcripts, potentially associated with anti-parasitic immunity, as well as increasing immunosuppressive molecules like PDL-1.

**Figure 1.**
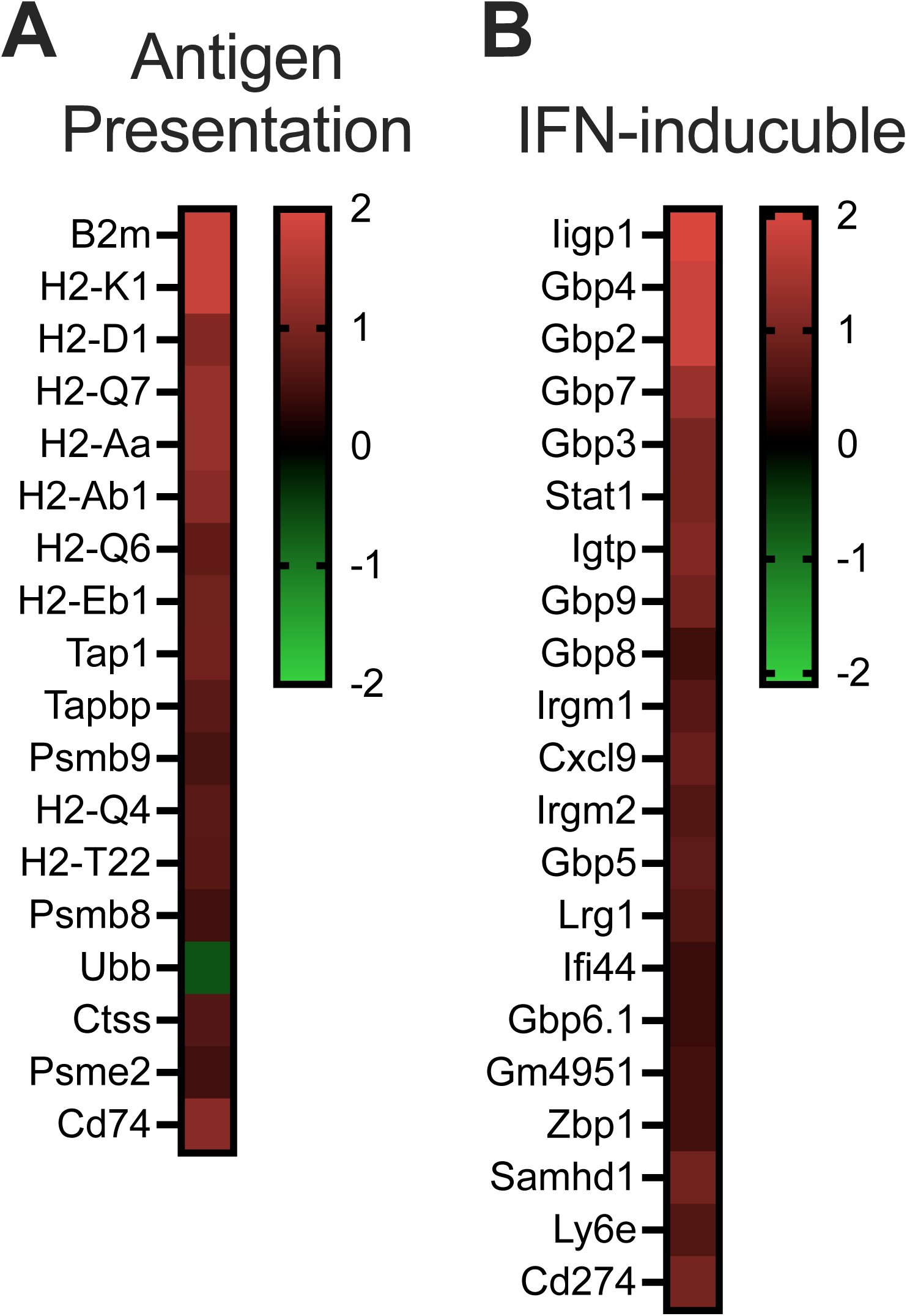
Single cell RNASequencing shows LEC activation in the lesion during *L. major* infection. C57BL/6 mice were infected with *L. major* parasites intradermally in the ear. At 4 weeks infected ears and naïve uninfected control ears were digested and subjected to scRNASequencing as a part of a previous study (20). Deeper analysis of the LEC cluster showed 114 transcripts with a significant adjusted p-value of <0.05. Within those top transcripts, 47 are upregulated with infection in LECs compared to LECs from naïve controls. The majority of the 47 upregulated transcripts are associated with antigen presentation (**A**) or have been identified as an IFN-inducible gene (IRG) (**B**). The color intensity represents the degree of expression. A red-green color scale was used to reflect the standardized gene expression with red representing high expression and green representing low expression.

Transcriptomic findings from scRNASeq suggest LECs are becoming activated with infection and undergoing translation to participate in the immune response to *L. major* parasites (20). For instance, transcripts of guanylate binding proteins (GBPs), MHC and PDL/PDL-1 molecules were elevated in LECs with infection (20). To determine the translational activity of LECs in vivo after infection, mice were infected, and at 5 weeks p.i. mice were injected with puromycin (32, 33). Puromycin incorporation as assessed by flow cytometry was used to measure the in vivo translation activity of dermal LECs (Fig. 2A). The translational activity of dermal LECs is significantly higher with *L. major* infection compared to LECs from the contralateral ear of the same mouse (Fig. 2B-C). Interestingly, even though infection enhanced LEC translational activity, the basal level translational activity of LECs is significantly higher than many other cell types in uninflamed skin (Fig. 2C-D). Of all the cell types analyzed in the skin, LECs and macrophages had the highest translational activity both at the site of infection and in the uninflamed contralateral ear (Fig. 2C-D). In addition to LECs, the translational activity of Ly6C^+^ inflammatory monocytes, macrophages, and blood endothelial cells (BECs) was significantly increased at the site of infection compared to the uninflamed contralateral ear (Fig. 2C). Of note, T cells had low levels of translational activity; in particular, CD8^+^ T cells had low levels of translational activity that further significantly decreased with infection (Fig. 2C). Altogether, these results suggest that dermal LECs exhibit high translational activity that is further elevated at the site of infection.

**Figure 2.**
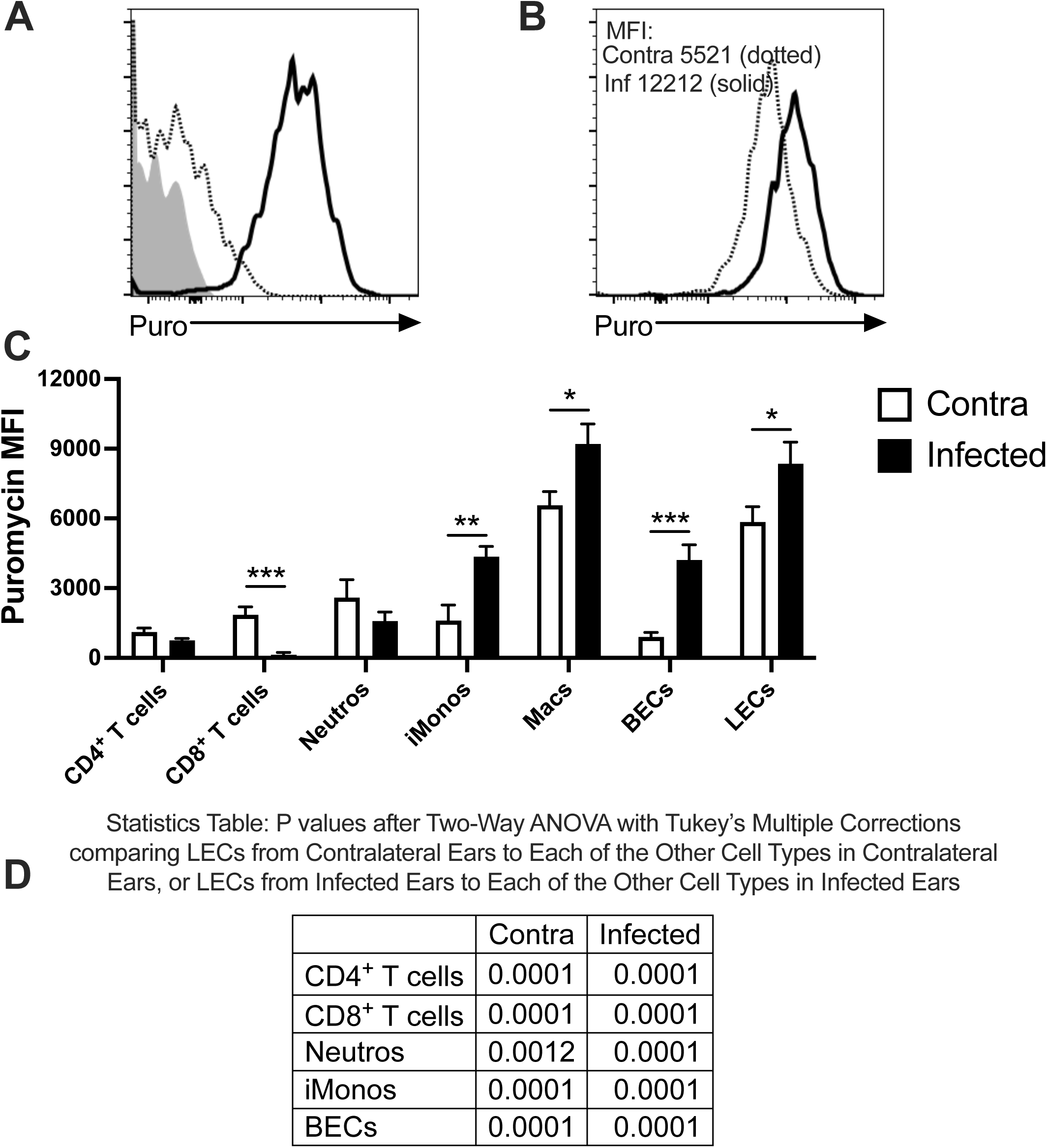
LECs exhibit increased translational activity following *L. major* infection. C57BL/6 mice were infected with *L. major* parasites intradermally in the ear. (**A**) Representative flow cytometry histogram plot showing controls for puromycin incorporation assay using LECs from an infected mouse. Mice were injected or not with puromycin (puro; 21.8 μg/g in PBS). Gated on live, single, CD45^-^CD31^+^podoplanin^+^ LECs. Grey = Fluorescence minus one (FMO) control for mouse injected with puromycin and stained for puromycin. Dotted line = Infected mouse that was not injected with puromycin but stained for puromycin. Solid line = puromycin-injected mouse and stained for puromycin. (**B-D**) At 3-5 weeks post-infection when mice exhibited lesions, mice were given puromycin i.p. 60 min before euthanizing. Infected and contralateral ears were processed, stained for puromycin, and analyzed by flow cytometry. (**B**) Representative flow cytometry histogram plot showing puromycin MFI after gating on CD45^-^CD31^+^podoplanin^+^ LECs from the infected or contralateral ear. (**C**) Quantification of puromycin MFI on endothelial and hematopoietic cell populations by flow cytometry including CD45^+^CD3^+^CD4^+^ T cells, CD45^+^CD3^+^CD8^+^ T cells, CD45^+^CD11b^+^Ly6G^+^ neutrophils, CD45^+^CD11b^+^Ly6G^-^Ly6C^+^ inflammatory monocytes, CD45^+^CD11b^+^Ly6G^-^CD64^+^ macrophages, CD45^-^CD31^+^podoplanin^-^ BECs, and CD45^-^ CD31^+^podoplanin^+^ LECs from infected or contralateral ears. All cells were pre-gated on total, live, singlets. Data shown here are pooled from two experiments with 10 mice per group total. Data are presented as the mean +SEM. *p < 0.05, **p < 0.01, ***p < 0.001, t-test comparing between infected and contralateral ears. (**D**) To compare LECs versus other cell types in leishmanial lesions, two-way ANOVA followed by the post hoc Tukey’s test comparing puromycin MFI between all cell types.

Transcriptomic findings from scRNASeq suggest LECs are participating in antigen presentation and processing. For instance, transcripts of MHC-I, MHC-II and immunoproteasome molecules were elevated in LECs with infection (20). To define the expression of MHC-II protein by LECs at the site of *L. major* infection, mice were infected, and at 5 weeks p.i., MHC-II expression was evaluated on LECs by flow cytometry. We found a significant increase in the percentage of LECs expressing MHC-II in the infected ear compared to the contralateral ear of the same mouse (Fig. 3A-B). In addition to MHC-II, LEC activation has been associated with elevated PDL-1 expression in melanoma, psoriasis, and viral infection (29, 40). Therefore, we evaluated PDL-1 on dermal LECs following *L. major* inoculation. At 5 weeks p.i., we discovered a significant increase in the percentage of LECs expressing PDL-1 in the infected ear compared to the contralateral ear of the same mouse (Fig. 3C-D). Taken together, these findings validate our previous scRNASeq results, and demonstrate that dermal LECs are activated in the skin following *L. major* infection.

**Figure 3.**
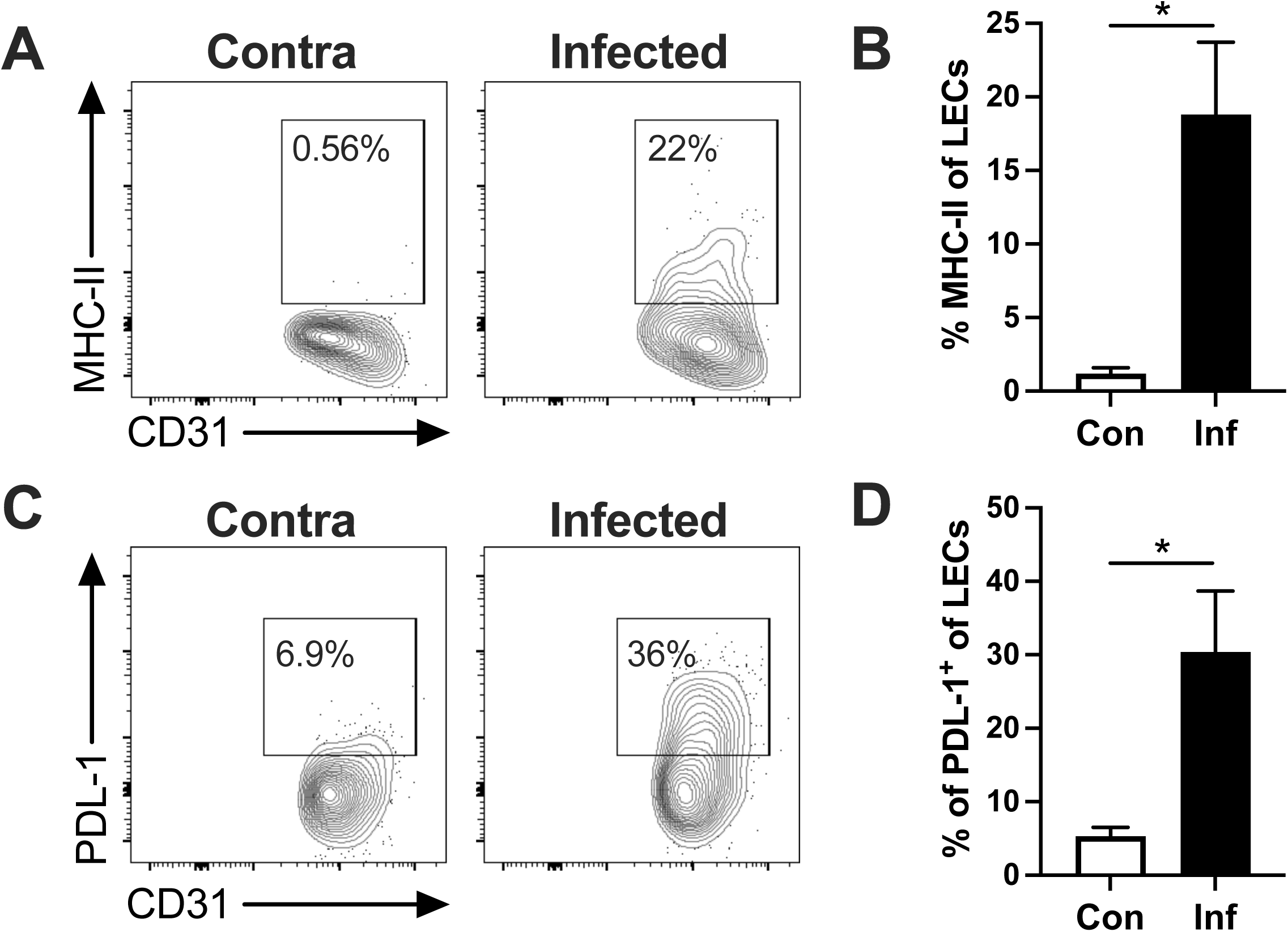
LECs express elevated MHC-II and PDL-1 during *L. major* infection. C57BL/6 mice were infected with *L. major* parasites intradermally in the ear. At 3-5 weeks p.i., MHC-II and PDL-1 expression were evaluated on LECs by flow cytometry. (**A**) Representative flow cytometry contour plots showing the percentage of dermal LECs expressing MHC-II protein after gating on CD45^-^CD31^+^podoplanin^+^ LECs from the infected or contralateral ear. (**B**) Quantification of the percentage of LECs expressing MHC-II at the site of *L. major* infection in the skin. (**C**) Representative flow cytometry contour plots showing the percentage of dermal LECs expressing PDL-1 protein after gating on CD45^-^CD31^+^podoplanin^+^ LECs from the infected or contralateral ear. (**D**) Quantification of the percentage of LECs expressing PDL-1 at the site of *L. major* infection in the skin. All cells were pre-gated on total, live, singlets. Data shown here is a representative experiment of 2-3 individual experiments with at least 4 mice per group. Data are presented as the mean +SEM. *p < 0.05, t-test comparing between infected and contralateral ears.

To determine if the activated LECs were also the same cells that exhibited increased translational activity, we evaluated activated LECs for translational activity using PDL-1 as a surrogate for LEC activation. To examine the translational activity, we gated on LECs that were positive or negative for PDL-1 (Fig. 4A), and evaluated the puromycin incorporation in these two subpopulations of LECs (PDL-1^+^ and PDL-1^-^ LECs). In a paired analysis, we found PDL-1^+^ LECs possessed a significantly higher puromycin median fluorescent intensity (MFI) compared to PDL-1^-^ LECs from the same *L. major*-infected mouse ear (Fig. 4B-C). This finding suggests increased translational activity is linked to LEC activation following *L. major* infection.

**Figure 4.**
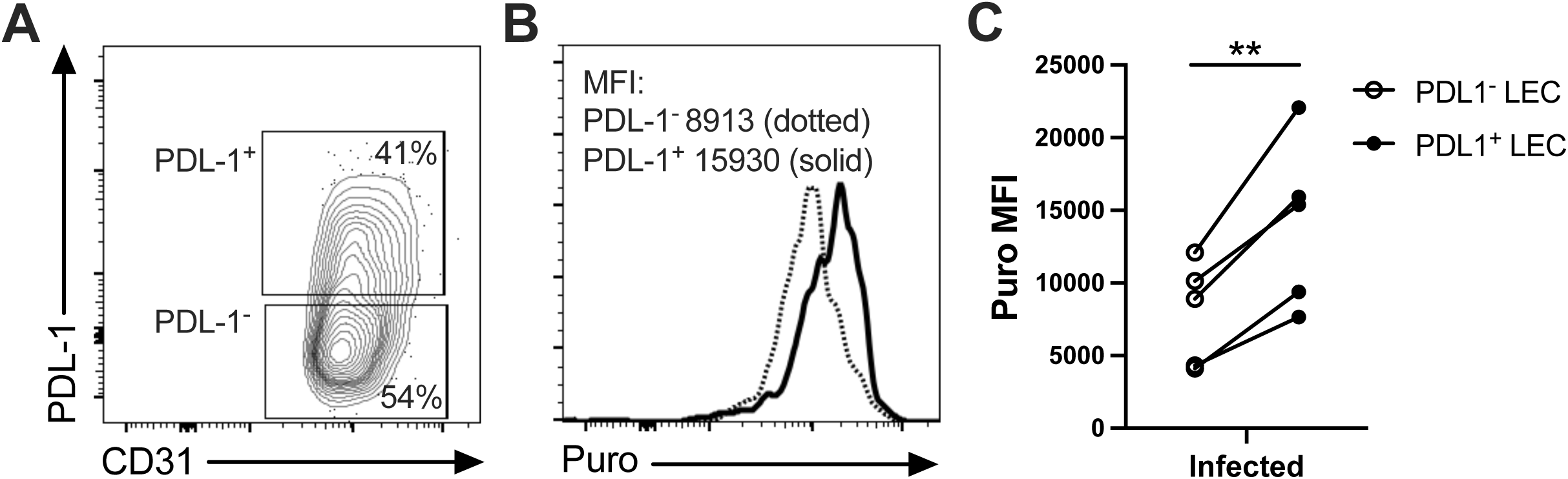
LEC activation is associated with increased translational activity during *L. major* infection. C57BL/6 mice were infected with *L. major* parasites intradermally in the ear. At 3-5 weeks p.i., mice were given puromycin and PDL-1 expression was evaluated on LECs by flow cytometry. (**A**) Representative flow cytometry contour plots showing subpopulations of dermal LECs that are PDL-1^+^ or PDL-1^-^ after gating on CD45^-^CD31^+^podoplanin^+^ LECs from an infected ear. (**B**) Representative flow cytometry histogram plot showing puromycin MFI after gating on PDL-1^+^ LECs or PDL-1^-^ LECs from an infected ear. (**C**) Quantification of paired puromycin MFIs from the same mouse comparing between PDL-1^+^ LECs and PDL-1^-^ LECs in the skin after *L. major* infection. All cells were pre-gated on total, live, singlets. Data shown here is a representative experiment of 2 individual experiments with at least 5 mice per group. **p < 0.01, paired t-test comparing PDL-1^+^ LECs and PDL-1^-^ LECs.

Based on the increase in MHCII^+^ LECs following infection, we determined the capacity of LECs to uptake and process antigen using DQ-OVA. Mice were infected, and at 5 weeks p.i. ear cells (single cell suspension) were placed in culture ex vivo with DQ-OVA, which resists acidic degradation and assesses antigen processing at 2 hours. LECs were identified by gating after flow cytometric analysis. To confirm the technical approach, we first examined CD11b^+^ dendritic cells (DCs), a major professional APC in leishmanial lesions that also takes up DQ-OVA (41). We detected a population of CD11b^+^ DCs that was positive for DQ-OVA after 2 hours ex vivo, confirming this assay assesses APC antigen uptake and processing using cells from leishmanial lesions ex vivo (Fig. 5A-B). After 2 hours, the percentage of LECs processing DQ-OVA was also significantly higher from the infected ears compared to LECs from the contralateral ears (Fig. 5C-D). Together, these data suggest LECs from the inflamed site of infection are significantly more capable in taking up and processing antigen than LECs from the uninflamed skin.

**Figure 5.**
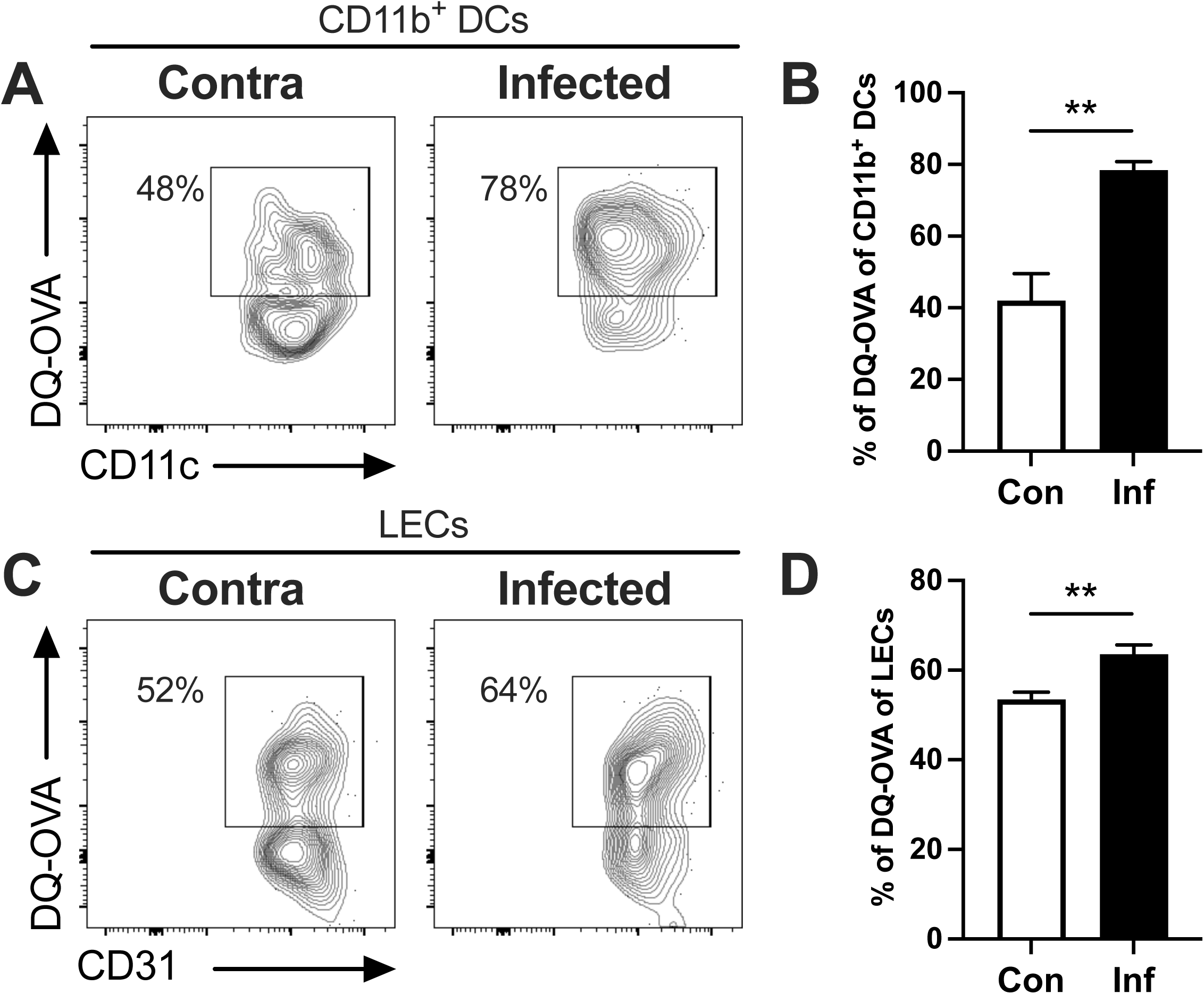
LECs from *L. major*-infected ears process antigen ex vivo. C57BL/6 mice were infected with *L. major* parasites intradermally in the ear. At 3-5 weeks p.i., digested ear cells were exposed to DQ-OVA for 2 hours in culture and evaluated by flow cytometry. (**A**) Representative flow cytometry contour plots showing the percentage of dermal CD11b^+^ DCs processing DQ-OVA after gating on CD45^+^CD11c^+^MHC-II^+^CD11b^+^ DC from the infected or contralateral ear. (**B**) Quantification of the percentage of dermal CD11b^+^ DCs from the site of *L. major* infection that are processing DQ-OVA. (**C**) Representative flow cytometry contour plots showing the percentage of dermal LECs processing DQ-OVA after gating on CD45^-^CD31^+^podoplanin^+^ LECs from the infected or contralateral ear. (**D**) Quantification of the percentage of dermal LECs from the site of *L. major* infection that are processing DQ-OVA. All cells were pre-gated on total, live, singlets. Data shown here is a representative experiment of 2 individual experiments with 5 mice per group. Data are presented as the mean +SEM. **p < 0.01, t-test comparing between infected and contralateral ears.

The DQ-OVA studies presented in Figure 5 are examining LEC antigen processing by skin cells ex vivo. However, the ability of LECs to process and present antigen in vivo during *L. major* infection is not known. To address this question, we injected DQ-OVA directly into leishmanial lesions to assess which cells, including LECs, could access and process antigen in their physiologic location in the inflamed dermal microenvironment. To confirm the technical approach, we first examined CD11b^+^ DCs, a major professional APC in lesions. At 5 weeks p.i., we detected a population of CD11b^+^ DCs that was positive for DQ-OVA in lesions within 24 hours after DQ-OVA injection, confirming this assay assesses APC antigen uptake and processing in vivo in leishmanial lesions (Fig. 6A-B). Similarly, we detected a population of LECs that was positive for DQ-OVA in lesions suggesting LECs can serve as APCs in vivo in leishmanial lesions (Fig. 6C-D). It should be noted that the LECs are negative for CD11b^+^ and CD11c^+^ and the LECs are gated from singlets (excluding doublets), suggesting the DQ-OVA detected on LECs is not due to professional APCs or myeloid cells that can be intimately associated with the lymphatic vasculature (42). Altogether, these results show that dermal LECs possess antigen presentation machinery and that these cells can uptake and process antigen during *L. major* infection.

**Figure 6.**
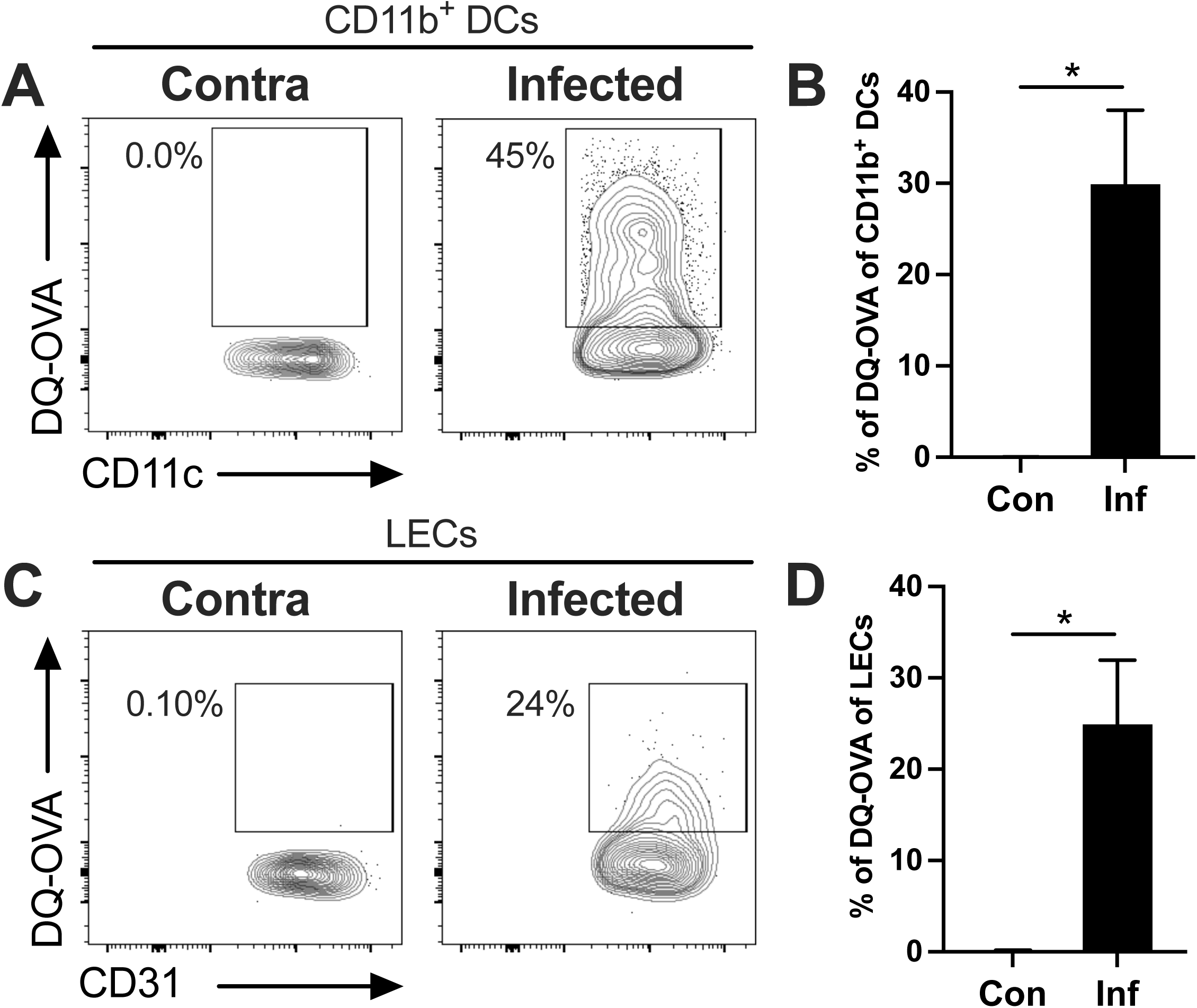
LECs process antigen in vivo during *L. major* infection. C57BL/6 mice were infected with *L. major* parasites intradermally in the ear. At 3-5 weeks p.i., DQ-OVA was injected into lesions. After 24 hours, digested ear cells were evaluated by flow cytometry. (**A**) Representative flow cytometry contour plots showing the percentage of dermal CD11b^+^ DCs processing DQ-OVA after gating on CD45^+^CD11c^+^MHC-II^+^CD11b^+^ DC from the infected or contralateral ear. (**B**) Quantification of the percentage of dermal CD11b^+^ DCs at the site of *L. major* infection that are processing DQ-OVA. (**C**) Representative flow cytometry contour plots showing the percentage of dermal LECs processing DQ-OVA after gating on CD45^-^CD31^+^podoplanin^+^ LECs at the infected or contralateral ear. (**D**) Quantification of the percentage of dermal LECs at the site of *L. major* infection that are processing DQ-OVA. All cells were pre-gated on total, live, singlets. Data shown here is a representative experiment of 2 individual experiments with 5 mice per group. Data are presented as the mean +SEM. *p < 0.05, t-test comparing between infected and contralateral ears.

## DISCUSSION

In CL, lesion severity results from uncontrolled parasite proliferation or an overactive host inflammatory response that impedes the healing process. Factors such as excessive numbers of neutrophils or CD8^+^ T cells, prior or concurrent viral infections, or the microbiome can exacerbate CL immunopathology, thereby increasing lesion size without altering parasite burdens (43–47). Cumulatively, the body of work from our lab indicates that the lymphatic vasculature also plays a role in immunopathology in CL (13–15). Specifically, we find lymphangiogenesis mitigates CL severity (13–15). Building on this foundation, here we find that LECs in leishmanial lesions become activated, increase translational activity, and enhance the ability to present antigens. These findings suggest LECs in leishmanial lesions might participate in the anti-parasitic immune response.

The role of LECs in immunity across inflammatory conditions is an area of active investigation. Currently, the prevailing view is that LECs are immunomodulatory, but most of those studies were performed using tumor models which are characterized by a immunomodulatory microenvironment. Therefore, it remains unclear whether LECs can promote pro-inflammatory immune responses. Our data suggests that in a robust pro-inflammatory microenvironment that is dominated by a Th1 immune response, such as a leishmanial lesion, LECs become activated exhibiting a mixed pro-inflammatory and immunomodulatory phenotype. Because LECs are not infected with parasites (20), the LEC activation is thought to be a result of the skin inflammatory microenvironment, rather than due to LECs harboring intracellular parasites. In this inflamed microenvironment, macrophages are the predominant cell type infected; however, it is not known how LECs interact with infected macrophages or how much of the inflammatory response is regulated by LECs in lesions. Regardless, we predict our findings are relevant to a variety of settings dominated by skin inflammation, and not restricted to CL.

The dominant perspective of the field is that LECs exert immunomodulatory effects, but the mechanisms of immunomodulation are not completely understood. For instance, LECs can influence immune responses by regulating dendritic cell (DC) maturation (23). Specifically, LECs can suppress DC activation and function when DCs are matured with TNFα, but not LPS (23). This observation, along with our findings, suggest LEC-DC interactions at sites of microbial infection differ from those at tumor sites or in general inflammation without a microbial involvement. Additionally, LECs modulate immune responses through inhibitory co-stimulatory molecules like PDL-1. Because IFNγ upregulates PDL-1 on LECs in culture, it is likely that PDL-1 on LECs in infected skin is induced by IFNγ (29), particularly given IFNγ is highly elevated in leishmanial lesions (48, 49). Alternatively, type I IFNs like IFNα may be responsible as IFNα also upregulates PDL-1 on LECs in culture (50) and IFNα is elevated in leishmanial lesions (51). Regardless of the factors influencing PDL-1 expression on LECs, our findings indicate that activated LECs (using PDL-1 positivity as a surrogate for activation) are associated with enhanced translational activity. To our knowledge, this is the first time that LEC translational activity was investigated in vivo. It is noteworthy that LECs were identified as one of the most translationally active cells within leishmanial lesions, and even in the absence of inflammation, which were unexpected observations. Future work should focus on whether the enhanced translational activity observed in LECs compared to other cell types is restricted to the skin or if it also occurs in LECs at other barrier or anatomical sites. Additionally, future studies will aim to determine if the heightened translational activity in LECs is specific to *Leishmania* infection or if it is also detected in skin inflammation due to other causes.

Under steady-state, LECs lack MHC-II antigen presentation machinery (52). However, IFNγ (which is elevated in leishmanial lesions (53, 54)) induces MHC-II on LECs, albeit less than other non-professional APCs (55, 56). While our transcriptomic findings suggest at least some MHC-II on LECs is endogenously generated during infection, MHC-II molecules on LECs can also be acquired from hematopoietic cells (55). Therefore, it is not clear from our analysis what portion of the LECs expressing MHC-II protein is due to endogenous expression or acquired from immune cells. Our findings are consistent with a previous report showing that LECs have antigen presentation machinery and immunoproteasomes during inflammation, though their exact role in immunity is not completely clear (56).

Future work will explore whether LEC antigen presentation boosts the anti-parasitic pro-inflammatory response, or if LEC antigen presentation functions similarly to other skin conditions like melanoma or viral infection where LECs are predominantly immunomodulatory. In these instances, LECs limit anti-viral immunity and prevent immunopathology (40). In contrast to melanoma and viral infection, we hypothesize that the LECs are taking on a more pro-inflammatory phenotype based on elevated MHC and GBP molecule expression from this work combined with our scRNASeq transcriptomic analyses of LECs after *L. major* infection. However, the elevated PDL-1 on LECs in CL also hints LECs may be involved in some immunomodulation of the T cell response during *L. major* infection. Given this discrepancy, a future objective of the lab is to determine whether LECs play a role in promoting anti-parasitic pro-inflammatory immune responses, or if they remain primarily immunomodulatory, as they have been in other conditions. This clarification will be crucial in understanding the role of LECs in immune responses.

### Limitations

Here, we compared the transcriptomic profiles of LECs from leishmanial lesions to uninfected uninflamed ears. To our knowledge this is the first comparison of this cell type following *Leishmania* infection in vivo. While these findings potentially shed light on LEC activation, there are limitations to these results. For instance, this work was only performed in a resistant C57BL/6 mouse model, and it is not obvious if the results will be replicated in a susceptible BALB/c mouse infected with *L. major* parasites.

Dermal LECs were also only assessed during a specific phase of *L. major* infection from 3-5 weeks p.i. when C57BL/6 mice exhibit lesions. However, over the course of infection, parasite numbers and lesion severity changes with time generating a dynamic tissue microenvironment. In resistant C57BL/6 mice, parasites are controlled, and lesions change over time with the inflammatory response lessening as the wound healing response takes over. As the lesion changes over time, LECs also respond to soluble mediators in the tissue microenvironment such as proinflammatory cytokines and growth factors as well as hypoxia. As a result, LECs in the tissue may exhibit variable activation states at different time points following infection depending on factors in the tissue microenvironment. Because lesions were mainly analyzed at 3-5 weeks p.i., it will be necessary to assess other time points following infection when LECs may exhibit altered activation states like 1-2 weeks p.i. when the IFNγ Th1 response is dominant but lesions are not present, 8-10 week p.i. when lesions are healing, and at 12 weeks p.i. when lesions have resolved.

## ACKNOWLEGMENTS/FUNDING STATEMENT

This work was supported by the Arkansas IDeA Network for Biomedical Research Excellence (INBRE) (funded by National Institutes of Health (NIH) National Institute of General Medical Sciences (NIGMS) Centers of Biomedical Research Excellence Grant P20-GM103429) and the Center for Microbial Pathogenesis and Host Inflammatory Responses (funded by NIH NIGMS Centers of Biomedical Research Excellence Grant P20-GM103625). This publication was also supported in part by funds provided by the National Center for Advancing Translational Sciences of the NIH under awards TL1 TR003109 and UL1 TR003107 for the Systems Pharmacology and Therapeutics (SPaT) NIH T32 training grant GM106999 to Lucy Fry. This work was also supported by the Arkansas INBRE PRO Program summer internship (funded by NIH NIGMS Grant P20-GM103429) to Alexx Weaver. This study was supported by the Arkansas Children’s Research Institute, the Arkansas Biosciences Institute, and the Center for Translational Pediatric Research funded under the National Institutes of Health National Institute of General Medical Sciences (NIH/NIGMS) grant P20-GM121293 and the National Science Foundation Award No. OIA-1946391. The content is solely the responsibility of the authors and does not necessarily represent the official views of the NIH. The funders had no role in study design, data analysis, decision to publish or preparation of the manuscript.

## AUTHOR DISCLOSURE STATEMENT

L. Fry: conceptualization, investigation, formal analysis, validation, visualization, review and editing. H. Roys: investigation, formal analysis, validation, review and editing. A. Bowlin: investigation, formal analysis, validation, review and editing. G. Venugopal: investigation, formal analysis, review and editing. J. Bird: formal analysis, visualization, review and editing. A. Weaver: investigation. S. Byrum: formal analysis, supervision, visualization, review and editing. T. Weinkopff: conceptualization, funding acquisition, investigation, formal analysis, supervision, validation, visualization, writing original draft and review and editing.

## COMPETING INTERESTS

The authors have declared that no competing interests exist.

## Notes

### Competing Interest Statement

The authors have declared no competing interest.

## REFERENCES

1. Alvar J, Velez ID, Bern C, Herrero M, Desjeux P, Cano J, et al. Leishmaniasis worldwide and global estimates of its incidence. PLoS One. 2012;7(5):e35671.

2. Rojas R, Valderrama L, Valderrama M, Varona MX, Ouellette M, Saravia NG. Resistance to antimony and treatment failure in human Leishmania (Viannia) infection. J Infect Dis. 2006;193(10):1375–83.

3. Leon B, Lopez-Bravo M, Ardavin C. Monocyte-derived dendritic cells formed at the infection site control the induction of protective T helper 1 responses against Leishmania. Immunity. 2007;26(4):519–31.

4. Petritus PM, Manzoni-de-Almeida D, Gimblet C, Gonzalez Lombana C, Scott P. Leishmania mexicana induces limited recruitment and activation of monocytes and monocyte-derived dendritic cells early during infection. PLoS Negl Trop Dis. 2012;6(10):e1858.

5. Ribeiro-Gomes FL, Peters NC, Debrabant A, Sacks DL. Efficient capture of infected neutrophils by dendritic cells in the skin inhibits the early anti-leishmania response. PLoS Pathog. 2012;8(2):e1002536.

6. Ribeiro-Gomes FL, Roma EH, Carneiro MB, Doria NA, Sacks DL, Peters NC. Site-dependent recruitment of inflammatory cells determines the effective dose of Leishmania major. Infect Immun. 2014;82(7):2713–27.

7. Bacellar O, Lessa H, Schriefer A, Machado P, Ribeiro de Jesus A, Dutra WO, et al. Up-regulation of Th1-type responses in mucosal leishmaniasis patients. Infect Immun. 2002;70(12):6734–40.

8. Bittencourt AL, Barral A. Evaluation of the histopathological classifications of American cutaneous and mucocutaneous leishmaniasis. Mem Inst Oswaldo Cruz. 1991;86(1):51–6.

9. Terabe M, Kuramochi T, Ito M, Hatabu T, Sanjoba C, Chang KP, et al. CD4(+) cells are indispensable for ulcer development in murine cutaneous leishmaniasis. Infect Immun. 2000;68(8):4574–7.

10. Gonzalez-Lombana C, Gimblet C, Bacellar O, Oliveira WW, Passos S, Carvalho LP, et al. IL-17 mediates immunopathology in the absence of IL-10 following Leishmania major infection. PLoS Pathog. 2013;9(3):e1003243.

11. Novais FO, Carvalho LP, Passos S, Roos DS, Carvalho EM, Scott P, et al. Genomic profiling of human Leishmania braziliensis lesions identifies transcriptional modules associated with cutaneous immunopathology. J Invest Dermatol. 2015;135(1):94–101.

12. Pereira LO, Moreira RB, de Oliveira MP, Reis SO, de Oliveira Neto MP, Pirmez C. Is Leishmania (Viannia) braziliensis parasite load associated with disease pathogenesis? Int J Infect Dis. 2017;57:132–7.

13. Bowlin A, Roys H, Wanjala H, Bettadapura M, Venugopal G, Surma J, et al. Hypoxia-Inducible Factor Signaling in Macrophages Promotes Lymphangiogenesis in Leishmania major Infection. Infect Immun. 2021;89(8):e0012421.

14. Weinkopff T, Roys H, Bowlin A, Scott P. Leishmania Infection Induces Macrophage Vascular Endothelial Growth Factor A Production in an ARNT/HIF-Dependent Manner. Infect Immun. 2019;87(11).

15. Weinkopff T, Konradt C, Christian DA, Discher DE, Hunter CA, Scott P. Leishmania major Infection-Induced VEGF-A/VEGFR-2 Signaling Promotes Lymphangiogenesis That Controls Disease. J Immunol. 2016;197(5):1823–31.

16. Fraga CA, Oliveira MV, Alves LR, Viana AG, Sousa AA, Carvalho SF, et al. Immunohistochemical profile of HIF-1alpha, VEGF-A, VEGFR2 and MMP9 proteins in tegumentary leishmaniasis. An Bras Dermatol. 2012;87(5):709–13.

17. Schatz V, Strussmann Y, Mahnke A, Schley G, Waldner M, Ritter U, et al. Myeloid Cell-Derived HIF-1alpha Promotes Control of Leishmania major. J Immunol. 2016;197(10):4034–41.

18. Araujo AP, Giorgio S. Immunohistochemical evidence of stress and inflammatory markers in mouse models of cutaneous leishmaniosis. Arch Dermatol Res. 2015;307(8):671–82.

19. Bowlin A, Roys H, Wanjala H, Bettadapura M, Venugopal G, Surma J, et al. Hypoxia inducible factor signaling in macrophages promotes lymphangiogenesis in Leishmania major infection. Infect Immun. 2021.

20. Venugopal G, Bird JT, Washam CL, Roys H, Bowlin A, Byrum SD, et al. In vivo transcriptional analysis of mice infected with Leishmania major unveils cellular heterogeneity and altered transcriptomic profiling at single-cell resolution. PLoS Negl Trop Dis. 2022;16(7):e0010518.

21. Viudez-Pareja C, Kreft E, Garcia-Caballero M. Immunomodulatory properties of the lymphatic endothelium in the tumor microenvironment. Front Immunol. 2023;14:1235812.

22. Card CM, Yu SS, Swartz MA. Emerging roles of lymphatic endothelium in regulating adaptive immunity. J Clin Invest. 2014;124(3):943–52.

23. Podgrabinska S, Kamalu O, Mayer L, Shimaoka M, Snoeck H, Randolph GJ, et al. Inflamed lymphatic endothelium suppresses dendritic cell maturation and function via Mac-1/ICAM-1-dependent mechanism. J Immunol. 2009;183(3):1767–79.

24. Montenegro-Navarro N, Garcia-Baez C, Garcia-Caballero M. Molecular and metabolic orchestration of the lymphatic vasculature in physiology and pathology. Nat Commun. 2023;14(1):8389.

25. Gkountidi AO, Garnier L, Dubrot J, Angelillo J, Harle G, Brighouse D, et al. MHC Class II Antigen Presentation by Lymphatic Endothelial Cells in Tumors Promotes Intratumoral Regulatory T cell-Suppressive Functions. Cancer Immunol Res. 2021;9(7):748–64.

26. Cohen JN, Guidi CJ, Tewalt EF, Qiao H, Rouhani SJ, Ruddell A, et al. Lymph node-resident lymphatic endothelial cells mediate peripheral tolerance via Aire-independent direct antigen presentation. J Exp Med. 2010;207(4):681–8.

27. Tewalt EF, Cohen JN, Rouhani SJ, Guidi CJ, Qiao H, Fahl SP, et al. Lymphatic endothelial cells induce tolerance via PD-L1 and lack of costimulation leading to high-level PD-1 expression on CD8 T cells. Blood. 2012;120(24):4772–82.

28. Hirosue S, Vokali E, Raghavan VR, Rincon-Restrepo M, Lund AW, Corthesy-Henrioud P, et al. Steady-state antigen scavenging, cross-presentation, and CD8+ T cell priming: a new role for lymphatic endothelial cells. J Immunol. 2014;192(11):5002–11.

29. Dieterich LC, Ikenberg K, Cetintas T, Kapaklikaya K, Hutmacher C, Detmar M. Tumor-Associated Lymphatic Vessels Upregulate PDL1 to Inhibit T-Cell Activation. Front Immunol. 2017;8:66.

30. Cohen JN, Tewalt EF, Rouhani SJ, Buonomo EL, Bruce AN, Xu X, et al. Tolerogenic properties of lymphatic endothelial cells are controlled by the lymph node microenvironment. PLoS One. 2014;9(2):e87740.

31. Spath GF, Beverley SM. A lipophosphoglycan-independent method for isolation of infective Leishmania metacyclic promastigotes by density gradient centrifugation. Exp Parasitol. 2001;99(2):97–103.

32. De Ponte Conti B, Miluzio A, Grassi F, Abrignani S, Biffo S, Ricciardi S. mTOR-dependent translation drives tumor infiltrating CD8(+) effector and CD4(+) Treg cells expansion. Elife. 2021;10.

33. Hidalgo San Jose L, Signer RAJ. Cell-type-specific quantification of protein synthesis in vivo. Nat Protoc. 2019;14(2):441–60.

34. Dobin A, Davis CA, Schlesinger F, Drenkow J, Zaleski C, Jha S, et al. STAR: ultrafast universal RNA-seq aligner. Bioinformatics. 2013;29(1):15–21.

35. Zheng GX, Terry JM, Belgrader P, Ryvkin P, Bent ZW, Wilson R, et al. Massively parallel digital transcriptional profiling of single cells. Nat Commun. 2017;8:14049.

36. Butler A, Hoffman P, Smibert P, Papalexi E, Satija R. Integrating single-cell transcriptomic data across different conditions, technologies, and species. Nat Biotechnol. 2018;36(5):411–20.

37. Stuart T, Butler A, Hoffman P, Hafemeister C, Papalexi E, Mauck WM, 3rd, et al. Comprehensive Integration of Single-Cell Data. Cell. 2019;177(7):1888–902 e21.

38. Macosko EZ, Basu A, Satija R, Nemesh J, Shekhar K, Goldman M, et al. Highly Parallel Genome-wide Expression Profiling of Individual Cells Using Nanoliter Droplets. Cell. 2015;161(5):1202–14.

39. McInnes L HJ, Melville J. UMAP: Uniform Manifold Approximation and Projection for Dimension Reduction. http://arxivorg/abs/180203426. 2018.

40. Lane RS, Femel J, Breazeale AP, Loo CP, Thibault G, Kaempf A, et al. IFNgamma-activated dermal lymphatic vessels inhibit cytotoxic T cells in melanoma and inflamed skin. J Exp Med. 2018;215(12):3057–74.

41. Gerner MY, Mescher MF. Antigen processing and MHC-II presentation by dermal and tumor-infiltrating dendritic cells. J Immunol. 2009;182(5):2726–37.

42. Cromer WE, Zawieja SD, Doersch KM, Stagg H, Hunter F, Tharakan B, et al. Burn Injury-Associated MHCII(+) Immune Cell Accumulation Around Lymphatic Vessels of the Mesentery and Increased Lymphatic Endothelial Permeability Are Blocked by Doxycycline Treatment. Lymphat Res Biol. 2018;16(1):56–64.

43. Novais FO, Carvalho AM, Clark ML, Carvalho LP, Beiting DP, Brodsky IE, et al. CD8+ T cell cytotoxicity mediates pathology in the skin by inflammasome activation and IL-1beta production. PLoS Pathog. 2017;13(2):e1006196.

44. Gimblet C, Meisel JS, Loesche MA, Cole SD, Horwinski J, Novais FO, et al. Cutaneous Leishmaniasis Induces a Transmissible Dysbiotic Skin Microbiota that Promotes Skin Inflammation. Cell Host Microbe. 2017;22(1):13–24 e4.

45. Crosby EJ, Clark M, Novais FO, Wherry EJ, Scott P. Lymphocytic Choriomeningitis Virus Expands a Population of NKG2D+CD8+ T Cells That Exacerbates Disease in Mice Coinfected with Leishmania major. J Immunol. 2015;195(7):3301–10.

46. Crosby EJ, Goldschmidt MH, Wherry EJ, Scott P. Engagement of NKG2D on bystander memory CD8 T cells promotes increased immunopathology following Leishmania major infection. PLoS Pathog. 2014;10(2):e1003970.

47. Santos Cda S, Boaventura V, Ribeiro Cardoso C, Tavares N, Lordelo MJ, Noronha A, et al. CD8(+) granzyme B(+)-mediated tissue injury vs. CD4(+)IFNgamma(+)-mediated parasite killing in human cutaneous leishmaniasis. J Invest Dermatol. 2013;133(6):1533–40.

48. Farias Amorim C, F ON, Nguyen BT, Nascimento MT, Lago J, Lago AS, et al. Localized skin inflammation during cutaneous leishmaniasis drives a chronic, systemic IFN-gamma signature. PLoS Negl Trop Dis. 2021;15(4):e0009321.

49. Bogdan C. Macrophages as host, effector and immunoregulatory cells in leishmaniasis: Impact of tissue micro-environment and metabolism. Cytokine X. 2020;2(4):100041.

50. Lucas ED, Finlon JM, Burchill MA, McCarthy MK, Morrison TE, Colpitts TM, et al. Type 1 IFN and PD-L1 Coordinate Lymphatic Endothelial Cell Expansion and Contraction during an Inflammatory Immune Response. J Immunol. 2018;201(6):1735–47.

51. Diefenbach A, Schindler H, Donhauser N, Lorenz E, Laskay T, MacMicking J, et al. Type 1 interferon (IFNalpha/beta) and type 2 nitric oxide synthase regulate the innate immune response to a protozoan parasite. Immunity. 1998;8(1):77–87.

52. Rouhani SJ, Eccles JD, Riccardi P, Peske JD, Tewalt EF, Cohen JN, et al. Roles of lymphatic endothelial cells expressing peripheral tissue antigens in CD4 T-cell tolerance induction. Nat Commun. 2015;6:6771.

53. Novais FO, Wong AC, Villareal DO, Beiting DP, Scott P. CD8(+) T Cells Lack Local Signals To Produce IFN-gamma in the Skin during Leishmania Infection. J Immunol. 2018;200(5):1737–45.

54. Scott P, Novais FO. Cutaneous leishmaniasis: immune responses in protection and pathogenesis. Nat Rev Immunol. 2016;16(9):581–92.

55. Dubrot J, Duraes FV, Potin L, Capotosti F, Brighouse D, Suter T, et al. Lymph node stromal cells acquire peptide-MHCII complexes from dendritic cells and induce antigen-specific CD4(+) T cell tolerance. J Exp Med. 2014;211(6):1153–66.

56. Santambrogio L, Berendam SJ, Engelhard VH. The Antigen Processing and Presentation Machinery in Lymphatic Endothelial Cells. Front Immunol. 2019;10:1033.

